# Event-Related Spatio-Spectral Perturbation (ERSSP): A Scalp-Wide EEG Analysis Framework for Capturing Extralemniscal Thalamocortical Activity, Validated Using a Fragile X Syndrome Model

**DOI:** 10.1101/2025.05.23.655841

**Authors:** Hyeonseok Kim, Lauren E. Ethridge, Lisa A. de Stefano, Lauren M. Schmitt, Ernest V. Pedapati, Craig A. Erickson, Makoto Miyakoshi

**Author notes:** **Corresponding author:** Makoto Miyakoshi. **Competing interests** The authors report no competing interests.

## Abstract

The extralemniscal thalamocortical pathway regulates large-scale modulation of cortical states but remains inaccessible to traditional EEG analysis due to its diffuse, variable, and asynchronous nature. To address this, we introduce the Event-Related Spatio-Spectral Perturbation (ERSSP) framework, which quantifies spatially distributed time-frequency responses. ERSSP summarizes the number of channels exhibiting significant spectral perturbations, capturing diffuse cortical dynamics from sensor-space EEG. We validated this approach using a passive auditory habituation task in individuals with Fragile X Syndrome (FXS), a condition marked by disrupted thalamocortical modulation. ERSSP revealed enhanced spatial extent of power increases (but not phase-locking) in FXS compared to controls. Results also showed expected gender differences, further supporting biological sensitivity of ERSSP. ERSSP provides a conceptually simple and computationally efficient method for quantifying broadly distributed sensory modulation and may enable new insights into thalamocortical function in both basic and clinical neuroscience.

## 1. Introduction

The human thalamus delivers information to the cortex through two partially independent channels. The lemniscal pathway starts from specific parts of the thalamus that receive direct sensory input — for example, the lateral geniculate nucleus for vision (1) and the medial geniculate body for hearing (2). Neurons in these regions send signals to layer IV of the primary sensory cortex, where they produce fast, focused brain responses that show up as early components in EEG recordings.

The extralemniscal pathway originates from thalamic regions known as the midline and intralaminar nuclei. Neurons in these areas send out widely branching axons that travel through the outermost layer of the cortex (layer I). These signals do not convey precise sensory content; instead, they regulate brain activity across large cortical areas, including the frontal, parietal, temporal, and cingulate cortices, supporting large-scale coordination by modulating cognitive states such as attention, arousal, and executive control (3). These broadly distributed inputs arrive later than core sensory signals and show delayed onset and contribute to sustained modulatory effects on cortical activity (4). They are thought to modulate cortical excitability (5) and influence stimulus salience (6), rather than encoding fine-grained sensory detail. Disruptions in thalamic matrix pathways are associated with sensory hyper-reactivity and vigilance regulation issues, and they have been observed in conditions like autism spectrum disorder (7), PTSD (8), and fragile X syndrome (FXS) (9).

Despite its relevance, extralemniscal function remains largely inaccessible to routine EEG analysis. Conventional event-related potential (ERP) approaches focus on a small number of midline electrodes, such as Cz, under the assumption that the response of interest will peak at a predictable scalp location. For example, ERP studies often quantify the N1 component at Cz or FCz to investigate auditory responses, including sensory habituation in clinical populations such as FXS (10). While such approaches have yielded insights into group differences in early sensory processing, they are inherently limited in their ability to capture the broad spatial patterns that define extralemniscal projections. Some studies have attempted to move beyond single-channel analysis using multivariate techniques such as principal component analysis (PCA), which combines signals across the scalp to maximize sensitivity to components like N1. For example, PCA has been applied to investigate reduced habituation in FXS (11). Similarly, independent component analysis (ICA) has been applied to isolate putative neural sources (12). Although these approaches differ in their mathematical foundations, with PCA maximizing shared variance and ICA separating statistically independent sources, they both rely on assumptions that make them poorly suited to capture extralemniscal activity. Specifically, both methods prioritize components that are expressed consistently across space and time. In contrast, extralemniscal signals tend to be spatially diffuse and less stereotyped in their temporal dynamics. They may influence many channels in a variable or asynchronous manner, which can limit the likelihood that such activity forms a dominant or clearly separable component. As a result, extralemniscal signals may be fragmented, misattributed, or entirely overlooked in PCA or ICA decompositions.

To address these limitations, we developed the Event-Related Spatio-Spectral Perturbation (ERSSP) framework, a sensor-space analysis strategy designed to capture the broad distribution of event-related spectral activity across the scalp. Unlike traditional ERP or component-based methods, ERSSP does not rely on inverse modeling, fixed electrode selection or assumptions about focal sources. Instead, it summarizes the number of channels showing significant spectral perturbations at each time–frequency point, providing a direct measure of spatial extent over time. This structure makes ERSSP well suited for detecting activity patterns associated with diffuse thalamocortical systems, including those arising from extralemniscal pathways. The framework is conceptually simple, computationally lightweight, and applicable at the level of individual subjects.

In this study, we introduce the ERSSP framework as a tool for quantifying extralemniscal activity from scalp EEG. To validate this approach, we applied it to high-density recordings collected during a passive auditory habituation task in individuals with FXS and typically developing controls. This paradigm offers a strong validation setting for ERSSP because auditory habituation engages thalamocortical circuits, and individuals with FXS consistently exhibit enhanced N1 responses and reduced habituation. These patterns are associated with dysregulated sensory gating and have been linked to dysfunction in thalamocortical systems, including extralemniscal pathways. By testing ERSSP under these conditions, we assessed whether the framework captures physiologically meaningful patterns linked to diffuse sensory modulation.

## 2. Methods and Materials

### 2.1 Dataset

We analyzed forty-seven individuals with Fragile⍰X⍰Syndrome (FXS; 19 females, 28 males) and 45 typically developing controls (TDC; 22 females, 23 males). The FXS group had a mean age of 25.1⍰±⍰8.4⍰years, while the control group had a mean age of 31.5⍰±⍰13.5⍰years. All individuals in the FXS group had a clinically confirmed diagnosis of Fragile⍰X⍰Syndrome, based on molecular testing (full-mutation or mosaic presentations). All procedures were approved by the Cincinnati Children’s Hospital Medical Center Institutional Review Board, and written informed consent (or guardian consent with participant assent) was obtained from every participant.

### 2.2 Habituation task

Participants completed an auditory habituation paradigm consisting of 150 stimulus trains. Each train contained four identical 50⍰ms white noise bursts separated by a 500⍰ms inter-stimulus interval, and consecutive trains were separated by a 4⍰s inter-trial interval. Stimuli were presented at 651dB1SPL through headphones. Stimuli were presented at 65⍰dB⍰SPL through headphones. During acquisition, participants watched a silent movie. No behavioural response was required.

**Figure 1.**
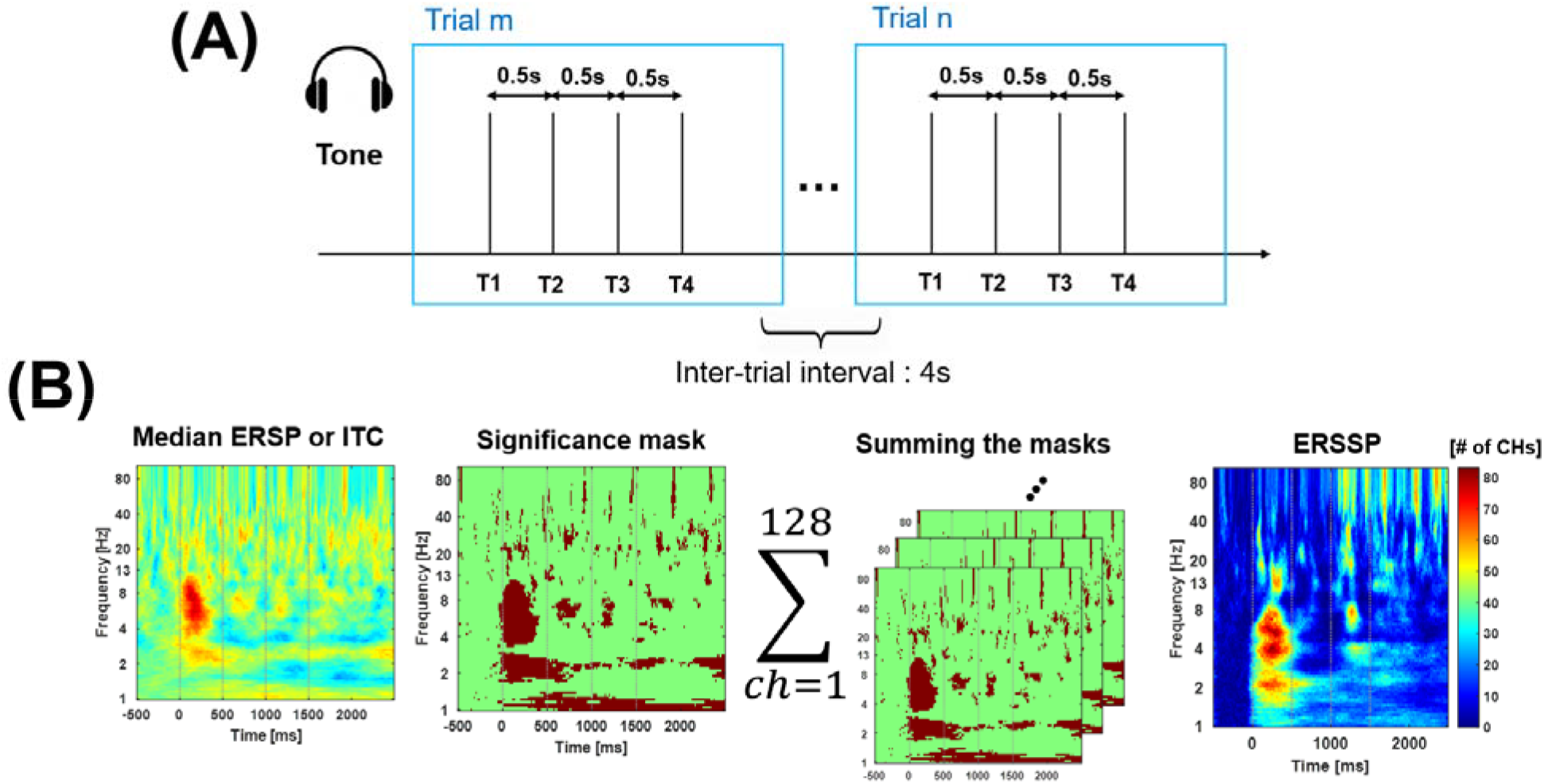
Overview of auditory habituation paradigm and ERSSP analysis pipeline. (A) Schematic of the auditory habituation task. Each trial consisted of four 50⍰ms white noise bursts (T1–T4), separated by 500⍰ms inter-stimulus intervals. Trials were presented every 4 seconds (inter-trial interval: 4⍰s), and a total of 150 stimulus trains were delivered at 65⍰dB SPL while participants watched a silent movie. No behavioral response was required. (B) Overview of the Event-Related Spatio-Spectral Perturbation (ERSSP) framework. Continuous EEG signals were first transformed using Morlet wavelets to produce time-frequency representations. For each subject, median ERSP or ITC values were computed across trials at each time-frequency bin (leftmost plot). Significant deviations from the baseline period were identified by thresholding each bin based on the cumulative distribution of baseline values (Significance mask). Binary masks from all 128 channels were then summed to yield a spatiotemporal map reflecting the number of channels with significant activity at each bin (Summing the masks). The resulting channel-count heatmap (rightmost plot) summarizes the broad spatial and spectral distribution.

### 2.3 EEG⍰Acquisition and Preprocessing

EEG data was collected with a 128-channel EGI system, sampled at 11kHz. All channels were referenced to the EGI system’s vertex reference electrode (located near the central midline), and recorded with a 0.01–200⍰Hz hardware band-pass filter. Electrode locations follow the International 10/10 system for 42⍰% of sensors, with a further 42⍰% positioned within 2⍰cm of their 10/10 analogues. EEG preprocessing was carried out in EEGLAB⍰ (v2024.1) (13). After data import, signals were resampled to 250⍰Hz, and the first and last 3⍰s of each recording were discarded. Continuous data were then high-pass filtered at 0.5⍰Hz (Blackman window, 1⍰Hz transition band) and low-pass filtered at 52.5⍰Hz (Blackman window, 5⍰Hz transition band) to attenuate high-frequency noise, including 60⍰Hz line noise present in the recording environment. Channel quality was assessed in two stages. First, putative flat-line channels were identified based on the standard deviation (SD) of each channel’s signal. Channels were marked as flat if their SD fell below a robust threshold defined as median(channelSD) – 4 × MAD(channelSD), where MAD is the median absolute deviation. If this threshold was less than or equal to 0, a fixed threshold of 0.1⍰µV was used instead, as a heuristic lower bound. Flagged channels were removed from the working montage. The data were then temporarily re-referenced to the median potential and segmented into 1,000 contiguous epochs. For each channel, the standard deviation was computed within each epoch, and the median of these values was used to assess channel-wise variability. Channels were marked as high-amplitude outliers if their median SD exceeded a robust threshold defined as the group-level median plus 20 times the scaled median absolute deviation (MAD × 1.4826). These outlier channels were marked for later interpolation. Artifact Subspace Reconstruction (ASR) was then applied with a burst cutoff of⍰25. ASR was calibrated using time windows where no channels were excessively noisy, effectively excluding windows with any bad channels from the calibration process (i.e., ref_maxbadchannels set to 0) (14). Channels flagged in the quality checks were subsequently restored by spherical spline interpolation, and the data were re-referenced to the average (including the original reference) (15). Finally, epochs from –0.5⍰s to⍰+2.5⍰s relative to the first burst of each four-burst train (t⍰= ⍰0) were extracted for all subsequent analyses.

### 2.4 ERP analysis

For the cleaned EEG that was segmented into epochs that began 500⍰ms before the first burst of each four-burst train and ended 2.5⍰s after that burst, The –500⍰to⍰0⍰ms interval served as baseline. Event-related potentials were measured at channel 55, which is located near the standard 10–20 Cz position in the 128-channel montage. For each of the four tones in the train, the N1 component was defined as the most negative-going peak between 60 and 140⍰ms post-stimulus. Peak amplitudes and latencies were identified for each stimulus repetition. To identify and exclude noisy trials within each subject, three robust criteria were applied to the target electrode. First, the standard deviation during the baseline period was computed for each trial, with high values reflecting baseline instability; trials were rejected if this value exceeded a threshold defined as the median plus 2 × scaled MAD (MAD × 1.4826). Second, the maximum amplitude across the entire epoch was assessed. This criterion was designed to detect abnormally large positive deflections inconsistent with typical post-stimulus ERP components. Third, the minimum amplitude was evaluated to capture extreme negative deflections that could mask or distort expected early components such as N1. Trials were rejected if either the maximum or minimum amplitude exceeded ±2 × scaled MAD from the median. Any trial exceeding any one of these criteria was excluded from further analysis. To ensure sufficient signal quality, subjects were excluded if their N1 signal-to-noise ratio (SNR) was below a minimum threshold. SNR was defined as the ratio of the N1 peak amplitude (relative to baseline mean, assumed to be 0) to the standard deviation of the baseline period. Specifically, SNR = N1_peak / baseline_SD, where baseline_SD is computed over the pre-stimulus baseline window. Subjects with an SNR less than 3 were considered to have insufficient ERP clarity and were excluded from further analysis. Habituation was quantified as the percentage change in N1 amplitude from the first tone to subsequent tones within each train. A repeated-measures ANOVA was conducted on percent change values, with Repetition (1st, 2nd, 3rd tone) as a within-subject factor and Group (FXS vs. control) as a between-subjects factor.

### 2.5 ERSSP framework

The continuous, artifact-clean signal was first transformed with a continuous wavelet transform (CWT) using Morlet wavelets. After the transform, the data were cut into the same –0.5⍰s to⍰+2.5⍰s epochs used for the ERP analysis; power values were converted to decibels with respect to the –0.5⍰s to⍰0⍰s baseline of each epoch, and inter-trial coherence (ITC) was calculated in parallel. To summarize each subject’s spectral response, the median power and ITC values across trials were calculated at each time-frequency bin. To capture the broad scalp distribution, a defining feature of extralemniscal activity, event-related spectral perturbation (ERSP) data from each channel were converted into binary significance masks on a subject-by-subject basis. For each subject and channel, ERSP values were statistically evaluated at every time-frequency point by computing their cumulative probability under a normal distribution derived from the baseline period’s mean and standard deviation. Bins falling in the top or bottom 5% of this distribution were marked as significant. Positive deviations indicated event-related synchronization (ERS), while negative deviations indicated desynchronization (ERD). The same procedure was applied to ITC, which reflects phase consistency across trials and ranges from 0 to 1. Since only increases in ITC are meaningful, bins in the top 5% relative to the baseline distribution were marked as significant. Time-frequency regions of interest (ROIs) were then defined based on the spatial extent of significant activity across the scalp. For each subject, binary significance masks were summed across channels to produce a time-frequency matrix indicating the number of channels with significant effects at each bin. These matrices were averaged across subjects, and bins exceeding the 95th percentile of the resulting matrix were grouped into spatially contiguous clusters. A rectangular ROI was defined to enclose the frequency and time span of the largest cluster. Any portion of the ROI extending into the pre-stimulus baseline period was excluded. To quantify extralemniscal activity, the average value within the previously defined time-frequency ROI was computed for each subject using the channel-summed significance matrix (either from ERSP or ITC), yielding a summary measure of the spatiotemporal extent of significant activity, referred to as Event-Related Spatio-Spectral Perturbation.

### 2.6 Validation

To evaluate the construct validity of the ERSSP as a measure of extralemniscal activity, we conducted a two-way ANOVA with group (FXS vs. TDC) and gender (male vs. female) as between-subjects factors. The dependent variable was the ERSSP value, computed separately from ERSP and ITC data (i.e., ERSSP-ERSP and ERSSP-ITC), and defined as the average number of significant time-frequency-channel bins within a group-defined ROI.

This validation approach was grounded in the known neurobiology of FXS, which is characterized by disrupted thalamocortical processing, including abnormal modulation of sensory input via extralemniscal pathways. These disruptions are broadly distributed and have been linked to reduced sensory gating and impaired auditory habituation which are functional features associated with extralemniscal thalamocortical circuits. FXS thus provides a clinically well-characterized model in which to test the sensitivity of ERSSP to biologically grounded dysfunction in this system A main effect of gender would reflect known sex-linked mechanisms in FXS, where males typically show greater disruption of sensory processing due to full mutation of the X-linked FMR1 gene, while females exhibit more variable expression due to mosaicism. Detecting such differences would support the sensitivity of the ERSSP metric to biologically grounded variation in extralemniscal pathway function. Together, these effects would support the interpretation of ERSSP as a biologically specific and generalizable marker of extralemniscal activity. All statistical analyses were conducted using MATLAB R2023b.

## 3. Results

To establish the validity of FXS as a model for evaluating extralemniscal sensory dysfunction, we first analyzed auditory ERPs, focusing on the N1 component and its habituation across stimulus repetitions. The repeated measures ANOVA revealed a significant main effect of Group, F(1, 72) = 8.67, p = .004, indicating reduced N1 habituation in the FXS group. There was also a significant main effect of repetition, F(2, 144) = 24.36, Greenhouse–Geisser corrected p < .0001, reflecting overall suppression of the N1 response across repeated tones. The group × repetition interaction was not significant, F(2, 144) = 0.23, p = .79. These results confirm a reduction in auditory habituation in individuals with FXS, replicating prior findings and supporting the use of this population and paradigm as a biologically grounded model for validating the ERSSP framework.

To assess distributed sensory responses potentially shaped by extralemniscal thalamocortical systems, we computed ERSSP values for each subject based on the number of scalp electrodes showing significant spectral perturbations within a group-defined time–frequency region of interest. ERSSP-ERS and ERSSP-ITC values reflect the spatial extent of significant power increases and phase consistency, respectively, and are expressed in units of channels. For ERSSP-ERS, a two-way ANOVA with Group (FXS vs. control) and Gender (female vs. male) as between-subjects factors revealed a significant main effect of group, F(1, 88) = 12.61, p = .00062, and a significant main effect of gender, F(1, 88) = 4.01, p = .048. The group × gender interaction was not significant, F(1, 88) = 0.87, p = .355. These results indicate that individuals with FXS exhibited greater spatial extent of spectral synchronization, consistent with diffuse cortical activation and extralemniscal system involvement. In contrast, ERSSP-ITC did not show any statistically significant effects. The main effects of group and gender were marginal (F(1, 88) = 3.07, p = .083; F(1, 88) = 2.99, p = .087, respectively), and the group × gender interaction was not significant, F(1, 88) = 0.73, p = .395. These results suggest that the spatial extent of event-related power increases, but not phase-locking, is altered in FXS. The ERSSP-ERS metric thus captures group-level differences that align with prior evidence of sensory processing abnormalities in FXS and offers a complementary perspective to traditional ERP-based measures.

**Figure 2.**
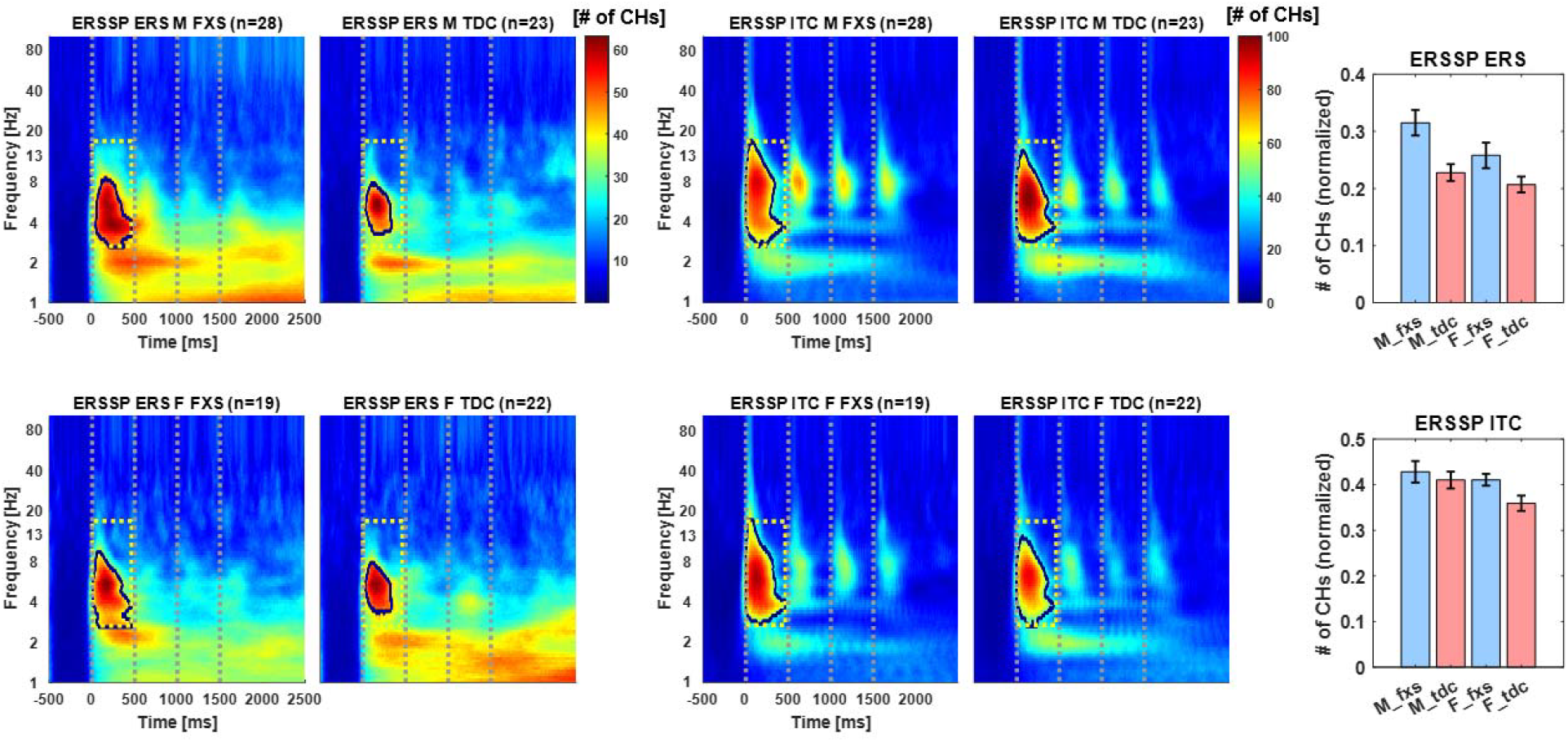
Group-level ERSSP maps and summary statistics. Time–frequency plots show the number of electrodes with significant event-related synchronization (ERS, left) and inter-trial coherence (ITC, middle) across male and female participants with Fragile X Syndrome (FXS) and typically developing controls (TDC). Warmer colors indicate more electrodes showing significant effects within a group-defined region of interest (dashed box). Bar plots (right) depict group means of normalized ERSSP values. FXS groups exhibited greater spatial extent of ERS compared to TDC, with significant main effects of group and gender; no significant effects were observed for ITC.

## 4. Discussion

In this study, we introduced the ERSSP framework as a new sensor-space method for detecting and characterizing extralemniscal thalamocortical activity from scalp EEG. Applying ERSSP to a well-established auditory habituation paradigm in individuals with FXS, we demonstrated that this metric reliably captures group-level differences in the spatial extent of event-related spectral power increases. Because ERSSP is designed to reflect broadly distributed cortical activity without relying on anatomical source modeling or inverse solutions, it is particularly well suited for investigating systems like the extralemniscal thalamocortical pathway. This makes it applicable in a wide range of contexts, including studies examining changes in arousal specifically at the beginning or end of the auditory stimulus presentation in an ASSR paradigm (Miyakoshi et al. 2025), as well as paradigms that manipulate inter-trial intervals to modulate salience responses via extralemniscal pathways (16). In clinical populations beyond FXS, such as individuals with autism spectrum disorder (17) or schizophrenia (18), ERSSP may support targeted investigations of thalamocortical dynamics by capturing the spatial extent of relevant activity directly from sensor-level data.

A notable advantage of the ERSSP framework lies in its implicit control over multiple comparisons. Because significance is assessed independently within each channel, and supra-threshold bins are binarized, the probability that multiple channels will simultaneously exceed threshold at the same time– frequency point under the null hypothesis is exceedingly low. This approach is consistent with the rationale used in spatial clustering-based analyses (19, 20), where random overlap across independent tests becomes increasingly improbable as the number of overlapping channels increases. As a result, ERSSP leverages the spatial independence of noise across electrodes, and the likelihood of false-positive convergence falls off approximately with p^k, where p is the per-channel threshold and k is the number of overlapping channels.

While we used a 95th percentile threshold for defining significant bins, the ERSSP framework is inherently flexible. Other thresholds, such as the 90th or 97.5th percentile, can be applied to match experimental needs or desired sensitivity. Similarly, although we defined ERSSP in terms of the raw number of supra-threshold channels, alternative scaling approaches are possible. For instance, z-scoring the channel-summed matrix or normalizing by subject-specific maximums could allow for cross-subject comparability in different populations. Nonetheless, expressing ERSSP in units of channels offers an intuitive interpretation aligned with the framework’s focus on spatial extent.

The current validation was conducted using high-density EEG. Future work is needed to assess how ERSSP performs when applied to recordings with lower channel counts, such as those used in clinical or portable systems. Additionally, while our findings support ERSSP’s sensitivity to extralemniscal activity, direct physiological validation using concurrent imaging remains an important next step. Despite these limitations, ERSSP provides a conceptually simple, robust, and scalable method for capturing distributed spectral dynamics that are otherwise inaccessible to traditional ERP or source-space analyses. The framework is openly available and may support novel insights in both basic and clinical neuroscience.

## References

1. Sherman SM (2007): The thalamus is more than just a relay. Curr. Opin. Neurobiol. 17(4): 417–22.

2. Jones EG (2001): The thalamic matrix and thalamocortical synchrony. Trends Neurosci. 24(10): 595–601.

3. Van der Werf YD, Witter MP, Groenewegen HJ (2002): The intralaminar and midline nuclei of the thalamus. anatomical and functional evidence for participation in processes of arousal and awareness. Brain Res. Brain Res. Rev. 39(2–3): 107–40.

4. Müller EJ, Munn B, Hearne LJ, Smith JB, Fulcher B, Arnatkevičiūtė A, et al. (2020): Core and matrix thalamic sub-populations relate to spatio-temporal cortical connectivity gradients. Neuroimage. 222: 117224.

5. Saalmann YB (2014): Intralaminar and medial thalamic influence on cortical synchrony, information transmission and cognition. Front. Syst. Neurosci. 8: 83.

6. Somervail R, Bufacchi RJ, Salvatori C, Neary-Zajiczek L, Guo Y, Novembre G, et al. (2022): Brain responses to surprising stimulus offsets: phenomenology and functional significance. Cereb. Cortex. 32(10): 2231–44.

7. Orekhova EV, Stroganova TA, Schneiderman JF, Lundström S, Riaz B, Sarovic D, et al. (2019): Neural gain control measured through cortical gamma oscillations is associated with sensory sensitivity. Hum. Brain Mapp. 40(5): 1583–93.

8. Harricharan S, Rabellino D, Frewen PA, Densmore M, Théberge J, McKinnon MC, et al. (2016): FMRI functional connectivity of the periaqueductal gray in ptsd and its dissociative subtype. Brain Behav. 6(12): e00579.

9. Razak KA, Binder DK, Ethell IM (2021): Neural correlates of auditory hypersensitivity in fragile x syndrome. Front. Psychiatry. 12: 720752.

10. Knoth IS, Lippé S (2012): Event-related potential alterations in fragile x syndrome. Front. Hum. Neurosci. 6: 264.

11. Ethridge LE, White SP, Mosconi MW, Wang J, Byerly MJ, Sweeney JA (2016): Reduced habituation of auditory evoked potentials indicate cortical hyper-excitability in fragile x syndrome. Transl. Psychiatry. 6(4): e787.

12. Makeig S, Westerfield M, Jung TP, Enghoff S, Townsend J, Courchesne E, et al. (2002): Dynamic brain sources of visual evoked responses. Science. 295(5555): 690–94.

13. Delorme A, Makeig S (2004): EEGLAB: an open source toolbox for analysis of single-trial eeg dynamics including independent component analysis. J. Neurosci. Methods. 134(1): 9–21.

14. Kim H, Chang C-Y, Kothe C, Iversen JR, Miyakoshi M (2025): Juggler’s asr: unpacking the principles of artifact subspace reconstruction for revision toward extreme mobi. J. Neurosci. Methods

15. Kim H, Luo J, Chu S, Cannard C, Hoffmann S, Miyakoshi M (2023): ICA’s bug: how ghost ics emerge from effective rank deficiency caused by eeg electrode interpolation and incorrect re-referencing. Front. Signal Process. 3:

16. Picton TW, Woods DL, Baribeau-Braun J, Healey TM (1976): Evoked potential audiometry. J. Otolaryngol. 6(2): 90–119.

17. Kamita MK, Silva LAF, Magliaro FCL, Fernandes FD, Matas CG (2021): Auditory event related potentials in children with autism spectrum disorder. Int. J. Pediatr. Otorhinolaryngol. 148: 110826.

18. Rosburg T, Boutros NN, Ford JM (2008): Reduced auditory evoked potential component n100 in schizophrenia--a critical review. Psychiatry Res. 161(3): 259–74.

19. Nichols TE, Holmes AP (2002): Nonparametric permutation tests for functional neuroimaging: a primer with examples. Hum. Brain Mapp. 15(1): 1–25.

20. Maris E, Oostenveld R (2007): Nonparametric statistical testing of eeg- and meg-data. J. Neurosci. Methods. 164(1): 177–90.

